# Y chromosome-coded HSATII repeats may contribute to higher incidence of cancer in men

**DOI:** 10.1101/2024.08.16.608341

**Authors:** Hedi Hegyi, Matej Lexa

## Abstract

It has been observed that men have a shorter lifespan and higher incidence of certain cancers than women. We postulate here that this phenomenon may be attributed to the “toxic Y chromosome”, owing to the expression of normally silent HSATII repeats in cancer, abundantly present on chromosome Y. Investigating the contribution of all repetitive elements to chromatin interactions using a genome-wide HiC set derived from a colon cancer cell line (HCT116), we found several anomalies regarding satellite HSATII elements when compared to other genomic repeat types: (i) HSATII repeats tend to form HiC pairs mostly with their own kind, and depleted in most heterologous repeat pairs, whereas other types of repeats readily form heterologous HiC pairs; (ii) when treated with 5aza, a chemotherapeutic agent, the number of long-distance HSATII-overlapping HiC pairs significantly decreased when compared to the untreated samples; (iii) and most importantly, in the treated samples interchromosomal HiC read pairs involving a chromosome Y-located repeat and an HSATII repeat on either end almost completely disappear. (iv) We also found that HSATII repeats lack overlap with other types of repeats.

To account for these anomalies we queried non-B DNA structure forming such as G-quadruplexes and transcription-related triplex forming. We found that Y chromosome-coded HSATII repeats form significantly stronger triplexes than HSATII repeats on other chromosomes, their strength significantly correlating with the number of HiC reads they overlap with. We surmise that the above phenomena may cause the higher incidence of cancer in men and may lead to the disappearance of the Y chromosome observed in cancerous cells in men.

## Introduction

It has been postulated that accumulation of repetitive elements on the Y chromosome lowers the survival of the XY heterogametic sex (i.e. males in mammalian species) dubbed the “toxic Y chromosome” hypothesis. According to the theory, the repeats become epigenetically silenced, however, Y heterochromatin can be lost through aging, which activates transposable elements and lowers male longevity by contributing to a higher incidence of certain cancers in men (Muyle et al., 2021).

In this work we analyzed the repeat content of data derived from a HiC-based study of HCT116 cells, a male colon cancer cell line (Spracklin et al.). HiC is a sequencing-based method used to characterize spatial proximity of DNA in the nucleus relying, as a rule, on Illumina pair-end sequencing. Because of the relatively short reads generated by this technology, traditional HiC analysis methods (Pal et al., 2019) generally ignore or fail to describe the repetitive fractions of genomes. Moreover, until recently, human reference genomes have not contained the most repetitive parts. Only lately, with improvements in long read sequencing technologies, did it become possible to assemble the human genome completely (CHM13 assembly)(Nurk et al., 2022; Rhie et al., 2023). This latest assembly additionally maps two thirds of the Y chromosome that were not available in previous assemblies. Also a number of large pericentromeric tandem arrays containing, among other sequences, large stretches of HSATII satellite repeats, were added to the CHM13 human reference genome.

Faced with the challenge of analyzing HiC data that would better represent the repetitive fraction we developed new tools (Lexa et al., 2022) and provided additional insights (Lexa and Hegyi, 2023) towards that goal. To look at possible associations between particular annotated repeats of the genome and HiC pairs mapping to them, we analyzed these overlaps between the annotations and HiC reads (represented as intervals). (So, e.g a HiC pair connecting two Alu repeats would be described as “Alu-overlapping read pair”.)

High-copy tandem satellites repeat families (alpha, beta, SATI, II, III) constitute 15% of the human genome (Hall et al., 2017). Many of these repeats are considered evolutionary relics with no assigned function. Perhaps the most prominent of these is the pericentromeric HSATII repeat, with a high number of copies on chromosomes 1, 2, 5, 6. 7, 10, 13, 14, 15, 17 21, 22 & Y (Hall et al., 2017). While HSATII repeats are transcriptionally silent in normal human tissue they do get transcribed into ncRNA in half of all human cancers (Younger and Rinn, 2015). Tanne et al. (2015) hypothesized that HSATII transcripts may have immunostimulatory properties and being normally silent, they are not under selective pressures to avoid immunostimulatory nucleotide motifs.

Jagannathan et al suggested that pericentromeric repeats do have a conserved function: they hold the chromosomes together by forming “chromocenters” (Jagannathan et al., 2018). HSATII repeats are not transcribed in normal human cells and their expression is associated with pathological phenomena, namely senescence and cancer (Miyata et al., 2021). The authors discovered that noncoding RNA (ncRNA) derived from HSATII repeats directly impairs the DNA binding of CTCF. They concluded that this CTCF disturbance increases the accessibility of chromatin and activates the transcription of SASP-like inflammatory genes, promoting malignant transformation (Miyata et al., 2021).

In this work we investigated the contribution of repetitive elements to genome-wide chromatin interactions using a HiC data set derived from cancerous tissue (Spracklin et al.) and found that HSATII has several idiosyncratic features, distinguishing it from other types of repeats. We found that HSATII repeats tend to form HiC pairs with their own kind, unlike most other types of repeats. This is in sharp contrast with other types of repeats that readily form heterologous HiC pairs with other types.

We also found that HSATII repeats have a tendency to form triplexes and investigated this feature for all chromosomes, and in more detail for chromosomes Y and 16. We found that chromosome Y has the strongest propensity to form triplexes when transcribed into RNA. We concluded that the triplex forming capability of the transcribed HSATII repeats may be relevant to their pathology.

## Results

After selecting 5 million reads from each of 6 samples of a Hi-C dataset derived from HCT116 cells, a colon cell line, (3 untreated, 2 treated with 5-aza, a demethylating agent, and one knock-out sample lacking methyltransferase activity) we submitted each to HiC-TE, a pipeline by Lexa et al. (2022) designed to analyze genomic repeats in HiC datasets.

The HiC-TE pipeline identifies HiC pairs where both reads of the HiC pair overlap with an annotated genomic repeat. The genomic repeats have been mapped previously by RepeatMasker (Smit et al., 2013-2015) to the most recent, T2T (“gapless”) human genome (Nurk et al., 2022). After completing HiC-TE for the six samples we removed those pairs that mapped to the same repeat copy, to avoid false positives (trivial pairing?).

For each set we counted the individual repeats and repeat pairs for each annotated repeat type and compared the counts to the expected numbers of pairs assuming a random pairing (for detail see HiC-TE manual at https://gitlab.fi.muni.cz/lexa/hic-te). Heatmaps were generated from the counts of the actual pairs divided by those of the expected pairs shown on a logarithmic scale (Fig 1) for SRR15458768 (Fig 1a), a treated and SRR15458782 (Fig 1b), an untreated sample.

**Fig. 1.**
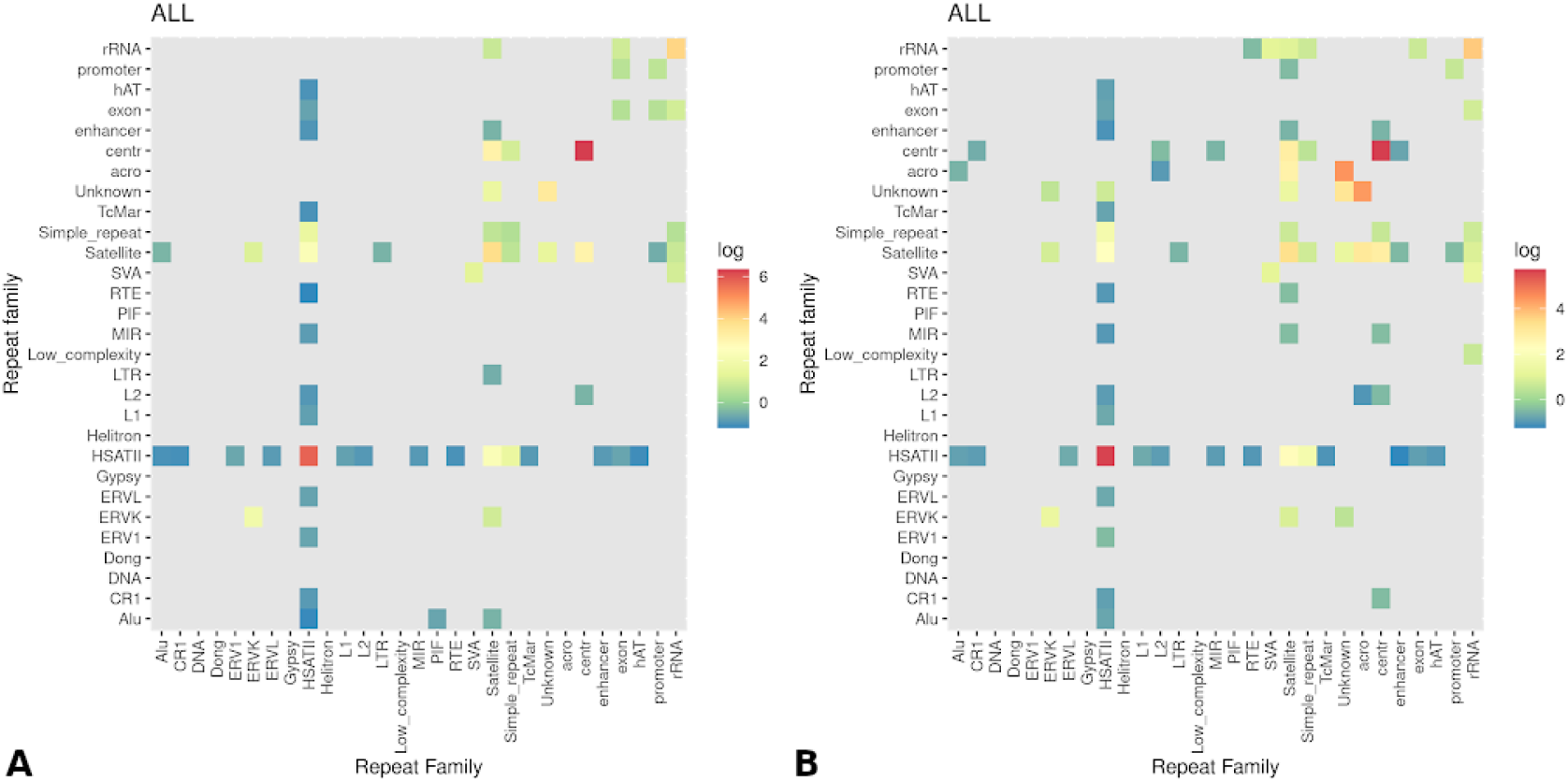
Actual/expected pair count ratios for each repeat pair in a (A) treated and a (B) untreated sample shown on a logarithmic scale.

Both the treated (Fig 1A) and untreated (Fig 1B) samples show the apparent tendency of HSATII repeats pairing with their own kind and being depleted in heterologous pairs. Repeat centr (RepeatMasker family centromeric satellite DNA) also tends to pair with its own kind, however, their absolute numbers are much lower than that of the HSATII repeats and are not known to be behind any known pathologies.

According to Bersani et al. (2015) HSATII RNA-derived DNA leads to progressive elongation of pericentromeric regions in tumors, where the reverse transcriptase activity is derived from human endogenous retroviruses (HERVs). To evaluate the possible involvement of HERVs in HSAT-related chromatin organization we looked at the ERVL-HSATII overlapping HiC pairs in an untreated (SRR15458780) and a treated (SRR15458784) sample using the ERVL type repeats of the annotated HERV set. The results are shown in Figure 2.

**Fig. 2.**
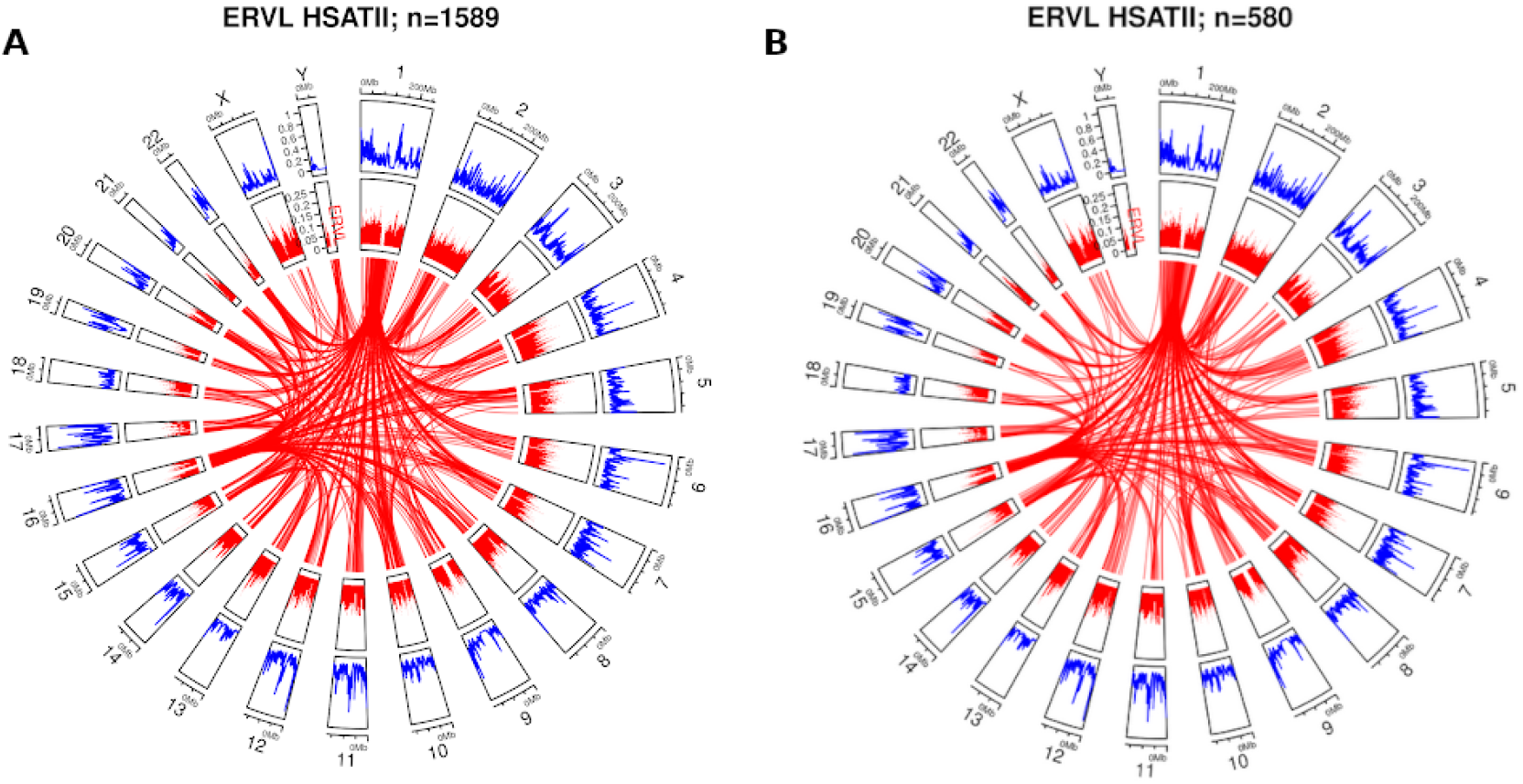
Chromosome-wise distribution of ERVL-HSATII overlapping HiC pairs in the (A) untreated SRR15458780 and (B) the treated SRR15458784 samples.

Apparently, the number of ERVL-HSATII pairs significantly decreased in the treated sample, supporting the observation of Bersani et al. (2015). Another interesting finding is the total lack of such pairs involving chromosome Y in the treated sample, a phenomenon we will further explore below.

In the next step we investigated repeat-overlapping HiC pairs when the two reads are on the same chromosome and one of the reads overlaps with an HSATII repeat. We counted the number of pairs within each distance range with a step of 100 kbp. We found that the distance distribution is markedly different for treated (Fig 3A) as opposed to untreated samples (Fig 3B) while the knockout (DKO) sample (Fig 3C) counts are somewhat between the treated and untreated sample counts. All counts for the 6 samples we studied are shown in Supplementary Table 1 (S1). A pairwise t-test was carried out for the 6 samples (Supplementary table S2) that showed that none of the treated samples differ significantly from each other or from the DKO sample but all three differ signifi-cantly from the untreated sample s80 shown in Fig 5B (with p-value < 9.9e-05).

**Fig. 3.**
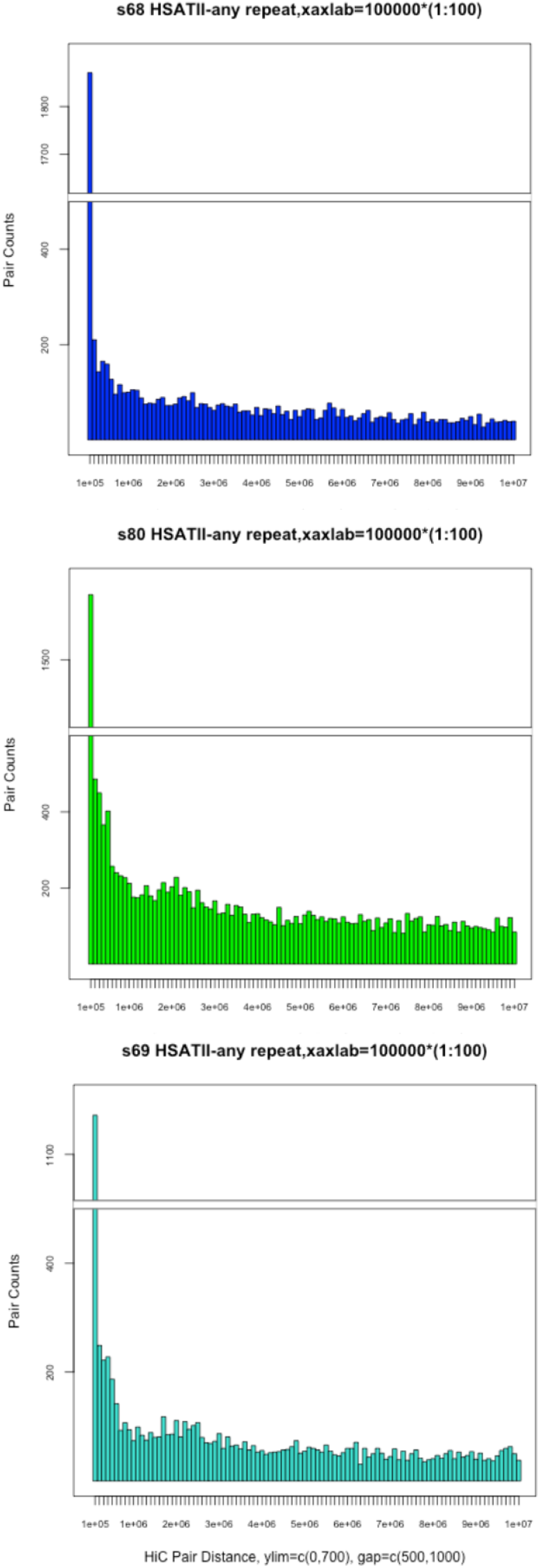
HiC pair distance distribution of pairs on the same chromosome where one of the pairs overlaps with a HSATII repeat. (A) Treated sample, SRR15458768; (B) Untreated sample SRR15458780; (C) Knockout (DKO) sample SRR15458769.

Overall, the untreated sample has more medium- (within a distance of 1Mb) and long distance (>1Mb) pairs where at least one of the reads overlaps with a HSATII repeat, short-distance pairs (within 100 kB) are more numerous in the treated sample (1871 vs 1671 pairs, Fig 3).

We also clustered the counts of the distances shown in Figure 3 for samples SRR15458768 and SRR15458780 for all the 6 samples. The resulting dendrogram is shown in Figure 4: while the two treated samples cluster with the knockout sample SRR15458769, the 3 untreated samples cluster with one another.

**Fig. 4.**
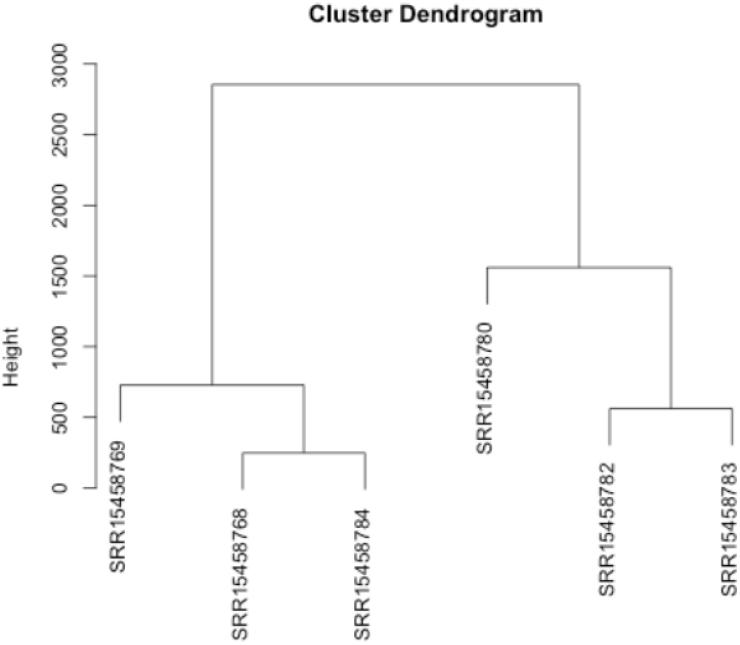
The clustering of the distance counts shown in Figure 3. Counts in the range of 0-20 million nucleotides were used based on the numbers in Supplementary table S1.

**Fig. 5.**
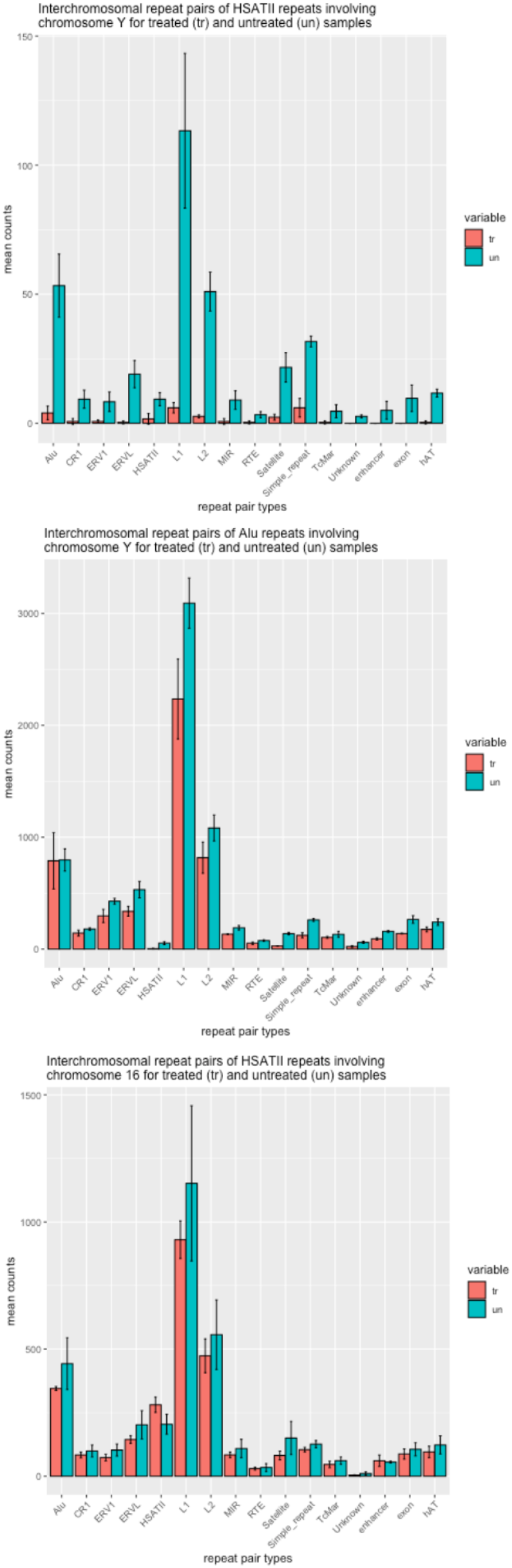
Interchromosomal repeat pairs of HSATII repeats involving chromosome Y for treated (tr) and untreated (un) samples. Error bars are SDs of the 3 counts in both groups.

We repeated the procedure of counting the intrachromosomal repeat pairs for each distance range similarly to the one shown in Figure 3 for all the other repeat types (where we required that one member of the pair overlaps with a particular repeat, while the other one overlaps with a repeat of any type). We found no significant difference between the treated, knock-out or untreated samples for any of the other repeat types (data not shown).

We also checked the interchromosomal HiC pairs counts, with one read overlapping a specific repeat type while its mate could be of any type (of repeat). While the overall statistics did not show a significant difference between the treated and untreated samples, unexpectedly there was a marked difference for pairs where one member overlaps a HSATII repeat and one member (either one) is located on the Y chromosome (Fig 5).

As shown in Fig 5A, the counts of basically all HSATII-containing repeat pairs with one of them located on chromosome Y are severely reduced in the treated samples when compared to the untreated ones. This is in sharp contrast with both other types of repeat pairs on chromosome Y (e.g. Alu-containing repeat pairs, Fig 5B) and HSATII-containing repeat pairs involving another chromosome (e.g. chr16-involving interchromosomal repeat pairs as in Fig 5C). This striking difference between treated and untreated samples involving specifically only chromosome Y and HSATII repeats strongly suggests their role in or contribution to chromosome Y-related pathologies (mostly cancer). The latter is consistently observed not only regarding colon cancer (such as in the current study) but for many other types of cancer as well (Miyata et al., 2023).

As non-B DNA structures (such as G-quadruplexes (G4s), triplexes, etc.) are known to be associated with many pathologies (for a review see (Wang and Vasquez, 2023)) we proceeded to check if HSATII repeats are associated with any of these structural variants. While we did not find any G4 enrichment for these repeats (data not shown), we did find a strong genome-wide association for RNA:DNA:DNA triplexes. We used PATO, an ultrafast triplex predictor (Amatria-Barral et al., 2023) using all genomic HSATII re-peats. The results for each chromosome are shown in Table 1.

**Table 1.** Triplex-forming score summaries and statistics using all genomic HSATII repeats summarized for each chromosome using PATO, a triplex predictor. In the last two columns the total length and the number of annotated HSATII repeats are shown. Chromosomes are ranked according to the PATO score.

| Chromosome | PATO score | PATO num | Total HSATII length | HSATII num |
| --- | --- | --- | --- | --- |
| chr 1 | 17 | 1 | 13364702 | 12013 |
| chr 4 | 23 | 1 | 6362 | 12 |
| chr 2 | 36 | 2 | 771636 | 168 |
| chr 20 | 44 | 2 | 19288 | 12 |
| chr 13 | 88 | 4 | 22203 | 53 |
| chr 14 | 113 | 5 | 30909 | 73 |
| chr 7 | 120 | 6 | 365335 | 95 |
| chr 9 | 126 | 6 | 165547 | 46 |
| chr 22 | 144 | 7 | 209195 | 107 |
| chr 21 | 277 | 13 | 90695 | 55 |
| chr 15 | 312 | 15 | 4722842 | 219 |
| chr 17 | 391 | 18 | 229286 | 43 |
| chr 10 | 473 | 20 | 648722 | 72 |
| chr 16 | 716 | 36 | 12848106 | 2413 |
| chr Y | 3616 | 168 | 8473577 | 6380 |

Remarkably, the highest triplex forming (single-stranded, i.e. assuming an RNA sequence) total score was reached for chromosome Y (3616), followed by chromosome 16 (716). While chr1 has the most HSATII repeats with the greatest genomic coverage (12013 repeats covering 13 Mbase), it has only one HSATII repeat predicted to form a triplex.

We also used TriplexAligner (Warwick et al., 2022), which predicts a score for each submitted sequence but it also requires a DNA decoy sequence. For the latter we used the longest transcript of the gene CTCF as hSATII RNA is known to change chromosomal interaction via CTCF disturbance Miyata et al. (2021).

Using TriplexAligner we predicted the strongest triplex forming between each HSATII repeat (covered by at least one HiC read in SRR15458782, an untreated sample) on chromosome Y and the longest transcript of CTCF. The top ten strongest triplexes are shown in Table 2A with the exact coordinates of the triplex in each HSATII repeat and also in the CTCF transcript (full list for each Y chromosome-coded HSATII is shown in Supplementary Table S3). We repeated the procedure for chromosome 16-coded HSATII repeats (also covered by a HiC read in SRR15458782), with the top 10 triplexes shown in Table 2B. Apparently, the Y chromosome-coded HSATII repeats form significantly stronger triplexes with scores 138.4-155.8 whereas the top chromosome 16 HSATII repeats fall into the 72.8-90.0 range.

**Table 2.** Top 10 predicted triplexes forming between the longest transcript of CTCF and HSATII repeats (coordinates shown) on (A) chromosome Y and (B) chromosome 16.

| HSATII repeat | Triplex start | Triplex end | CTCF transcript | Triplex start | Triplex end | Score [-log(e-value)] |
| --- | --- | --- | --- | --- | --- | --- |
| chrY:29076553-29078090 | 109 | 621 | chr16:73357234-73433945 | 351 | 863 | 138.470 |
| chrY:29002563-29004409 | 114 | 1847 | chr16:73357234-73433945 | 38747 | 39454 | 139.293 |
| chrY:28826798-28829040 | 706 | 1351 | chr16:73357234-73433945 | 262 | 907 | 139.464 |
| chrY:28838008-28841183 | 1020 | 1628 | chr16:73357234-73433945 | 309 | 917 | 140.426 |
| chrY:28974375-28977541 | 666 | 1469 | chr16:73357234-73433945 | 311 | 1114 | 140.975 |
| chrY:29169549-29173005 | 936 | 1535 | chr16:73357234-73433945 | 311 | 910 | 141.154 |
| chrY:29032431-29034017 | 651 | 1360 | chr16:73357234-73433945 | 311 | 1020 | 142.157 |
| chrY:61386197-61388802 | 1937 | 2335 | chr16:73357234-73433945 | 38748 | 39146 | 143.841 |
| chrY:61155405-61158796 | 2688 | 3391 | chr16:73357234-73433945 | 38747 | 39450 | 148.944 |
| chrY:61192479-61195935 | 1196 | 1910 | chr16:73357234-73433945 | 38747 | 39461 | 155.815 |

| HSATII repeat | Triplex start | Triplex end | CTCF transcript | Triplex start | Triplex end | Score [-log(e-value)] |
| --- | --- | --- | --- | --- | --- | --- |
| chr16:40283685-40286832 | 1551 | 1735 | chr16:73357234-73433945 | 18812 | 18996 | 72.812 |
| chr16:50701389-50718723 | 920 | 1099 | chr16:73357234-73433945 | 18817 | 18996 | 73.414 |
| chr16:51216503-51217806 | 920 | 1099 | chr16:73357234-73433945 | 18817 | 18996 | 73.414 |
| chr16:39595378-39601444 | 417 | 617 | chr16:73357234-73433945 | 18812 | 19012 | 73.849 |
| chr16:39680112-39683475 | 417 | 617 | chr16:73357234-73433945 | 18812 | 19012 | 73.849 |
| chr16:49644454-49652409 | 6 | 190 | chr16:73357234-73433945 | 18812 | 18996 | 73.944 |
| chr16:41704139-41722378 | 12590 | 12774 | chr16:73357234-73433945 | 18812 | 18996 | 74.435 |
| chr16:50421802-50428243 | 417 | 601 | chr16:73357234-73433945 | 18812 | 18996 | 75.016 |
| chr16:35672831-35674023 | 1078 | 1193 | chr16:73357234-73433945 | 38772 | 38887 | 80.196 |
| chr16:35653860-35663003 | 8983 | 9140 | chr16:73357234-73433945 | 38747 | 38904 | 90.032 |

To see if the triplex forming can be related to the HiC pair forming on chromosome Y we calculated the Pearson correlation between the number of HiC reads each HSATII repeat overlaps with and the strength of triplex forming using the TriplexAligner scores. To normalize the counts, instead of using the full-length triplex-forming ranges (as higher scores are usually associated with longer segments within the HSATII repeats), we used 2000-nucleotide ranges for each predicted triplex starting with the actual predicted 5’ location. We found that for the 3 untreated samples the Pearson R-values between the number of overlapping HiC reads and the triplex-forming score were 0.685, 0.426 and 0.531 (n=419, p-value < 1e-5 for all three samples). This latter result is in line with the finding of others (Jalali et al., 2017) who also found HiC pair enrichment around potential triplex-forming genomic locations.

## Discussion

While HSATII repeats are normally silent and part of the heterochromatin in normal cells, age-related senescence often causes these pericentromeric regions enriched in human satellite II (HSATII) to open up, dramatically changing the chromosomal conformation in these regions (Miyata et al., 2023), significantly contributing to cancer. In this study we show that HSATII repeats display a set of peculiar characteristics when studied by HiC, demethylation treatment and non-B DNA secondary structure sequence analysis. These characteristics are specific for HSATII and not seen in other repeat types annotated by Repeat Masker.

By analyzing 6 samples of Spracklin et al. that queried HCT116 cells-derived data using Hi-C technology, we found numerous anomalies uniquely associated with HSATII repeats, namely:

i. HSATII repeats tend to form Hi-C pairs with their own kind whereas most other repeats have no such preference (Fig 1), a phenomenon that may be related to their triplex-forming tendencies. As shown by Farabella et al. (2021) 3D genome organization is promoted by triplex-forming RNAs.
ii. HSATII-overlapping intrachromosomal Hi-C pair distance distributions markedly differ for treated (Fig 3A) and untreated (Fig 3B) samples where the longer-distance pairs are present in higher proportion in the untreated samples even at a genomic distance of 10 Mb. As shown by Taberlay et al. (2016) cancer cells organize their TADs (topologically associated domains) into smaller domains and the new boundaries coincide with genomic mutations with relevance to regulatory genes (e.g. TP53). The differences shown in Fig 3 are in line with this observation as short-distance HiC pairs do diminish in the untreated samples. Interestingly, no other repeat types show differences in their intrachromosomal distance distributions between the treated and untreated samples.
iii. The most striking differences appear for the HSATII-overlapping interchromosomal pair counts where one Hi-C read overlaps with a HSATII repeat and one (either one) is on chromosome Y. As shown in Fig 5A, the pair counts decrease very significantly in the treated samples whereas this is not the case for either different repeat types (Alu, Fig 5B) or other chromosomes (chr16, Fig 5C). As the treatments consisted of a demethylation agent, aza-5 and the related methyl transferase knockout, the disappearance of the chrY-related HSATII repeat overlapping contacts may clearly indicate the importance of methylation in maintaining proper chromatin organization and the possible Y chromosome involvement in HSATII biology.
iv. We also counted the fraction of repeats for all repeat types that overlap with other types based on the genomic annotations by RepeatMasker (Supplementary table S4). Remarkably, HSATII had the smallest fraction of its repeats overlapping with other types (12.4%) whereas most other repeat types overlapped to a considerably higher extent with a median value of 92.3% (Supplementary table S4). This is consistent with the fact that pericentromeric satellite repeats often form homogenous stretches of heterochromatin (Duggan and Tang, 2010) although the centromeric centr repeat type had a higher proportion of overlaps (52%) with other types of repeats. It may also indicate the unique sequences of HSATII repeats that are potentially toxic when expressed as RNAs as suggested by Hall et al. (2017)).
v. The propensity of chrY-coded HSATII repeats to form triplexes with the highest scores, as shown by both PATO and TriplexAligner, also support the notion that chromosome Y-coded HSATII are the hardest for the cellular mechanisms to resolve. As shown by Kaushal and Freudenreich (2019) triplexes above a certain length show replication fork stalling and cause chromosome fragility. This may lead to the loss of the Y chromosome (LoY) observed in many male cancers (Müller et al., 2023). The authors also note that LoY correlates with general genome-wide instability manifesting in frequent loss of autosomes (autosomal aneuploidy), usually preceded by LoY. As several human autosomes have a high number of HSATII repeats (listed in Table 1), our theory of demethylation of HSATII repeats as the cause of higher incidence of cancer and LoY would fit into this picture.

## Conclusions

Putting it all together we argue that the HSATII repeat-related anomalies as observed by studying HiC-derived samples from the HCT116 male cancer line and comparing them to treated samples may well account for cancer-related differences between men and women and consequently perhaps also for the shorter lifespan of men.

## Methods and data

After downloading 6 HiC datasets from the GEO website accompanying a recent publication by Spracklin et al. at https://www.ncbi.nlm.nih.gov/geo/query/acc.cgi?acc=GSE182105 (3 untreated samples [SRR15458780, SRR15458782, SRR15458783], from HCT116, a colon cell line, 1 sample [SRR15458769] a genetic derivative lacking DNMT3B and DNMT1 activity (referred to as DKO) and 2 samples [SRR15458768, SRR15458784] treated with the DNA methylation inhibitor 5-Azacytidine, a chemotherapeutic agent), we selected 5 million reads from each sample and submitted them to the HiC-TE pipeline (Lexa et al., 2022) designed to analyze genomic repeats in HiC datasets.

HiC-TE produces two kinds of tables, one, a BED formatted table of all HiC reads overlapping a genomic re-peat (with extension .bed) and one that contains all mapped HiC pairs where each read maps to a genomic repeat (with extension .tab) . For genomic repeats we used the annotated repeats downloaded from the UCSC genome browser that previously mapped all repeats to the T2T gapless human using RepeatMasker (https://pubmed.ncbi.nlm.nih.gov/19274634/). Both .bed and .tab files are uploaded to the zenodo site at https://shorturl.at/fmQbm

For Figure 1 we selected only those repeat-overlapping HiC pairs where the two reads of each HiC pair map to different repeats (we called clean set). The zipped script_all.zip (also available at the zenodo site, referenced above) file contains the shell script called remove_selfhics.sh that carries out this task. (All scripts used throughout the work are contained in this zip file.) To generate Figure 1 we used the generate_heatmap_reference_joint_probability_norm.R R script (part of the HiC-TE pipeline) on the clean set. To generate the chromosome-wise ERVL-HSATII repeat pair counts as shown in Figure 2 we used the plot_circos.nf workflow, which is also part of the HiC-TE pipeline.

To produce Figures 3A-C, we calculated the genomic distance for each HiC pair with the reads located on the same chromosome using the .tab output files of the HiC-TE pipeline. The length distribution figures (Figs 3A-C) were generated by R scripts called s68_gapbarplotHSATII.R, s80_gapbarplotHSATII.R and s69_gapbarplotHSATII.R, respectively.

The dendrogram shown in Figure 4 was produced with an R script called dendrogram_hclust.R using the histogram values shown in Fig 3 (but including all distance ranges whereas for Figs 3A-C we used distance values only up to 20 Mbase). After counting the interchromosomal HiC pairs separately for each of the 6 samples, where one of the overlapping repeats is an HSATII (or any other specific repeat type) and one of the pairs is on chrY, we calculated the mean and SD for each repeat pair type separately for the untreated and the treated samples (including the DKO sample in the latter). The results shown in Fig 5A were produced by an R script (Fig5a_chrY.R). By replacing HSATII with Alu we produced Fig 5B and replacing chrY with chr16 we produced Fig 5C. We also produced an R script (ttest_columns_selectrows_clever.R) carrying out a t-test between the two sets of samples and calculated the associated p-values for the HSATII-containing interchromosomal repeat pairs (chrY_reppairs_treated_vs_untreated_ttest_pvalue.tsv3).

We repeated the calculations for each of the other repeat types.

To generate all HSATII repeat sequences we used the bedtools getfasta command but first concatenated those repeat annotations (as identified by RepeatMasker) where two HSATII repeats coordinates overlapped. To carry out this task we used an iterative process written in perl called iterate_repeat_concat.pl that calls the bedtools intersect sub-command repeatedly till all overlapping repeat coordinates are expanded over the full length of the overlapping HSATII repeats. Once the HSATII repeat sequences were generated we ran the PATO software on them with the -ss switch, i.e. assuming single-stranded RNA sequences.

We used TriplexAligner to predict triplex forming between the longest transcript of the gene of CTCF and 419 HSATII repeats on chromosome Y that were selected based on the SRR15458782 sample having at least one HiC read mapping to these HSATII repeats. We selected the chr16 HSATII repeats using the same criteria (1784 HSATII repeats).

After predicting the TriplexAligner scores for chrY we took the BED output of the HiC-TE pipeline for each of the 3 untreated samples and intersected them (using the bedtools intersect subcommand with the -c switch to count the number of intersecting HiC reads) with the 419 HSATII repeats with predicted triplex scores. Instead of using the full predicted triplex-forming ranges (as determined by TriplexAligner) we used 2000 nucleotides-long ranges for each of the HSATII repeats, starting out at the actual 5’ end of the predicted triplex region. By using a uniform length for each predicted triplex region we can avoid length-based bias.

After counting the number of intersections between the HSATII-based triplexes on chrY and the HiC counts in each of the three untreated samples we used a Perl script to calculate the Pearson correlation R values and determined the corresponding p-values with the help of an online server at https://www.socscistatistics.com/pvalues/pearsondistribution.aspx.

## Supplementary data

Supplementary figures, tables, files, scripts and input data are available via a Zenodo repository at https://zenodo.org/uploads/13293981

